# The impact of transplant location on the gut microbiome and resistome in patients undergoing hematopoietic stem cell transplantation at home versus in the hospital

**DOI:** 10.1101/2024.11.19.624359

**Authors:** T.M. Andermann, K. Zeng, S. Guirales-Medrano, A. Groth, B.C. Ramachandran, S. Sun, A.A. Sorgen, L. Hill, A.T. Bush, H. Liu, C. Jones, J. Roach, B.P. Conlon, G. Rao, N.J. Chao, A.A. Fodor, A.D. Sung

## Abstract

**Objectives:** Home-based hematopoietic stem cell transplantation (HCT) is a novel approach that has the potential to improve outcomes, however, the impact of transplant location on the gut microbiome remains uncharacterized. We hypothesized that patients randomized to undergo home HCT would have higher gut taxonomic diversity and lower antimicrobial resistance (AMR) gene abundance compared to those undergoing standard hospital HCT.

**Methods:** We identified 28 patients enrolled in Phase II randomized trials of home (n=16) v. hospital (n=12) HCT at Duke and performed shotgun metagenomic sequencing of stools to compare taxonomic and AMR gene composition between groups. We performed a secondary analysis of patients from each group transplanted at an outpatient infusion clinic with those who underwent standard inpatient HCT (“outpatient” v. “inpatient”).

**Results:** No significant differences in duration of hospitalization were found in those randomized to home v. hospital HCT. Taxonomic and AMR gene α- and β-diversity were comparable. In contrast, secondary analyses demonstrated that patients from both home and hospital groups transplanted at an outpatient infusion clinic spent significantly less time in the hospital and demonstrated higher taxonomic α-diversity and differential β-diversity compared to standard inpatient HCT, although AMR gene α-diversity did not differ, and comparisons were confounded by both differences in transplant type and use of antibiotics.

**Conclusions:** Randomization by transplant location did not impact the gut microbiota to the same extent as the duration of hospitalization, although secondary analyses were heavily confounded. Even when taxonomic differences were observed, AMR genes were similar between groups.

## INTRODUCTION

Hematopoietic stem cell transplantation (HCT) is a potentially life-saving process that remains enormously burdensome for more than 22,000 individuals in the United States each year as of 2021^1^. Hospital stays for HCT are often several weeks to months in duration and result in significantly decreased quality of life^2–5^. Patients in the hospital are also at increased risk for exposure to multidrug-resistant nosocomial pathogens with each day spent as inpatient, through contact with healthcare personnel and the hospital environment(ref).

In contrast to hospital HCT, many programs offer outpatient HCT, in which patients receive all necessary peri-HCT care in an outpatient “day hospital” and return to home or temporary lodging at night. However, patients and their caregivers must still make daily trips to the day hospital, resulting in additional burden and strain. A new approach, home HCT, delivers all post-HCT care in the home setting, offering patients comfort through a familiar environment^2–5^. Patients in home HCT protocols in the U.S. and elsewhere generally receive pre-HCT conditioning and stem cell infusion in the hospital or day hospital, followed by immediate discharge home. Daily visits to the patient home (“house calls”) are performed by the healthcare team allowing for labs, vitals checks, and administration of medications. Prior case-controlled studies of home HCT in Sweden, for example, demonstrated lower rates of acute graft-versus-host disease (aGVHD), improved treatment-related mortality, and decreased medical costs^3,4,6,7^. The transplant group at Duke University demonstrated in a small Phase I observational study that, while clinical outcomes such as acute GVHD and infection did not differ between trial patients undergoing HCT at home versus standard of care hospital controls, pre-HCT quality of life was preserved for home HCT recipients^5^. Phase II randomized controlled trials are ongoing by this same group with patients accruing to trials investigating both autologous and allogeneic HCT at home (clinicaltrials.gov: NCT02218151 and NCT03667599). In these trials, researchers hypothesize that patients undergoing HCT at home are likely to have a reduced incidence of acute GVHD and infection risk in larger studies with more power to demonstrate differences between groups.

In patients enrolled and randomized to HCT at home versus in the hospital at Duke University, our group compared changes in the gut microbiome over time between these two groups. We hypothesized that patients undergoing HCT at home would have less disruption to their gut microbiomes over time, demonstrating greater gut microbial diversity following HCT relative to controls in the hospital. Knowing that the gut microbiome also serves as an important reservoir for multidrug-resistant organisms, we also asked whether patients undergoing HCT at home would demonstrate a decreased burden of antimicrobial resistance (AMR) in the organisms found in their gut.

## METHODS

### Patient cohort selection

Samples for the study were obtained retrospectively from a stool biospecimen repository of patients enrolled in the Phase I and Phase II trials of home versus hospital HCT at Duke between 2017 and 2020. Details for the Phase I and Phase II trials are provided elsewhere^5^. Research activities were supported by the Institutional Review Boards of Duke University (Duke IRB protocols #00032263, #00051024, and #00089697 under PI Dr. Anthony Sung, MD) and the University of North Carolina at Chapel Hill (UNC IRB protocol #20-2371 under PI Dr. Tessa Andermann, MD MPH). Stools in the biorepository had previously been collected at pre-defined time points (pre-HCT, and days 0, 7, 14, 21, 28, 35, 60, 100, 180, 270, 365, and 730 after HCT). Due to small sample size at D28 and D35, these were merged into a single bin labeled D30, referred as “day +30”, used in the majority of the analyses.

Patients were selected for inclusion if they had provided at least one stool sample prior to, and 2 samples following, either autologous or allogeneic HCT. Stool samples were excluded if there were fewer than 4 aliquots per patient available in the biobank. All stools available that met criteria from each patient were included in the study.

All allogeneic and autologous patients in both home and hospital HCT cohorts received standard antimicrobial prophylaxis with a fluoroquinolone--either ciprofloxacin or levofloxacin--along with acyclovir, fluconazole, and trimethoprim-sulfamethoxazole as previously described^5^.

Clinical metadata were collected prospectively from the electronic medical record and included date of transplantation, type of pre-HCT conditioning, donor type, graft source, duration of hospitalization, antimicrobial use, and date of aGVHD with aGVHD grade defined according to the IBMTR grading system^8^.

### Defining transplant location: inpatient versus outpatient HCT

In the Duke clinical trials, both autologous and allogeneic HCT recipients were randomized to either home or hospital HCT Patients randomized to either home or hospital HCT received their stem cell infusions either in the hospital (“inpatient”) or in the infusion clinic (“outpatient”) as decided upon by their transplant physician. We determined that inpatient and outpatient groups almost perfectly differentiated patients who were hospitalized for longer (inpatient) or shorter (outpatient) durations (see Supplementary Figure S4). For the purposes of investigating the impact of duration of hospitalization on the microbiome and resistome, we compared these two groups as part of our secondary analysis.

### Stool sample processing and sequencing

#### DNA extraction

Stools were collected from patients and placed on ice, stored at 4°C for less than 24 hours before aliquoting into cryovials, and frozen at -80°C before DNA extraction. DNA isolation was performed using an optimized version of the QIAamp Fast DNA Stool Mini Kit (Cat No./ID: 51604) protocol supplemented with 60 mg/mL lysozyme (Thermo Fisher Scientific, Grand Island, NY)^9^. At the time of DNA extraction, samples were transferred to a 2 ml tube containing 200 mg of 106/500μm glass beads (Sigma, St. Louis, MO) and 0.5 ml of Qiagen PM1 buffer (Valencia, CA). Mechanical lysis was performed for 40 minutes on a Digital Vortex Mixer. After a 5 minutes centrifugation, 0.45 ml of supernatants was aspirated and transferred to a new tube containing 0.15 ml of Qiagen IRS solution. The suspension was incubated at 4°C overnight. After a brief centrifugation, supernatant was aspirated and transferred to deep well plates containing 0.45 ml of Qiagen binding buffer supplemented with Qiagen ClearMag Beads. DNA was purified using the automated KingFisher™ Flex Purification System and eluted in DNase free water. ZymoBIOMICS Gut Microbiome Standard (Cat# D6331) was used as the positive control. Samples that had discolored DNA from the automated isolation process, underwent repeated DNA isolation via a column-based isolation protocol. Samples that had DNA concentration below 1 ng/uL from the automated isolation, also underwent repeated DNA extraction via column-based isolation.

#### Library preparation and metagenomic sequencing

DNA isolated from samples was assessed for sample quality and DNA fragment size on a TapeStation (Agilent, Santa Clara, CA). DNA libraries for sequencing were prepared using the Illumina DNA Seq protocol (formerly, Nextera Flex; Illumina Inc., San Diego CA). Samples were pooled in groups of 96 and then checked on a MiSeq Nano (Illumina Inc., San Diego CA) for pool balance. The pool was then rebalanced and sequenced as 150 paired-end reads on a NovaSeq 6000 S4 flowcell (Illumina Inc., San Diego CA). Sequencing output from the NovaSeq platform was converted to fastq format and demultiplexed using Illumina Bcl2Fastq 2.20.0.

#### Taxonomic and metabolic pathway characterization

Quality control of the demultiplexed sequencing reads was verified by FastQC 0.11.8^10^. Adapters were trimmed using TrimGalore 0.6.2^11^. The resulting paired-end reads were classified with Kraken 2.1.2^12^ and Bracken 2.5^13^ and all reads identified as host were eliminated. Reads unclassified by Kraken2 were submitted to Spades 3.14.1 for meta-assembly^14^. Contigs in the meta-assembly of length greater than 500 were submitted to BLAST 2.12.10^15^ for potential taxa identification. Reads were mapped back to an index of the contigs via Bowtie2 2.4.1^16^ for quantification. Contigs of the meta-assembly of reads identified as unknown were submitted to DFAST 1.2.15^17^ for annotation. Annotations were submitted to MetaQUAST 5.0.2^18^ for quality assessment. Taxa identified in the meta-assemblies of the unclassified reads were evaluated for suitability of inclusion in the Kraken2 database. Genomic references for suitable taxa were added from GenBank based on whether they were assembled, and at least specified a genus. This process continued for three iterations, at which point no suitable taxa were recovered. All reads identified as not being derived from host, including those that remained unclassified were submitted to Spades for meta-assembly. Contigs of these meta-assemblies were submitted to BLAST and Kraken2 for potential taxa identification and reads were mapped using Bowtie2 to the contigs for quantification. Annotation and quality assessment on the reads classified as non-host was performed using DFAST and MetaQUAST respectively. Potential metabolic pathways were characterized using Humann3^19,20^. Raw counts from the Bracken pipeline and pathway abundance were normalized using the formula:

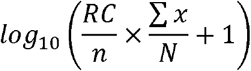

where RC represents the individual sample OTU counts, n is the number of sequences in a sample, the sum of x is the total number of counts in the table and N is the total number of samples.

To remove potentially spurious low abundance taxa, we removed all taxa or pathways with a relative abundance below 0.000168% at the species level, based on histogram distribution (data not shown) and corresponding to a mean relative abundance of 100 reads or less.

#### Antimicrobial resistance (AMR) gene characterization

Paired-end reads identified as non-host were joined with VSEARCH 1.17.1^21^ and queried against The Comprehensive Antibiotic Resistance Database (CARD)^22^. Non-host contigs assembled by Spades underwent antibiotic resistance gene (ARG) calling with CARD’s Resistance Gene Identifier 5.2.1 (RGI)^22^ and NCBI’s Antimicrobial Resistance Gene Finder 9.27 (AMRFinderPlus) in parallel^23^. RGI was run with default parameters which used a blast query algorithm to match genes in the CARD database to those in the contigs. RGI by default reports genes matched with 95% identity (“Strict”) up to 100% identity (“Perfect”). To assure the validity of the genes reported by RGI, AMRFinderPlus was run on the same contigs using default parameters. Genes reported from AMRFinderPlus were also restricted to genes that had EXACT (100% identity) and BLAST (>90% identity). AMR genes identified using each of these three methods were normalized using RPKM (reads per kilobase of transcript per million reads mapped). Genes with 80% or more of the samples with zero counts were filtered out.

### Statistical analysis

During each filtering step, i.e., raw reads, deduplicated reads, trimmed reads, and cleaned reads (host DNA removed), bbmap 38.96 was used to calculate read length. Read length mean and median summaries were calculated. To visualize patient sample relative to their collection date a time-scaled scatterplot was created. Taxonomic data from Bracken and antibiotic resistance gene data from each ARG caller were gathered and parsed into count tables.

Principal coordinate analysis (PCoA) or Multidimensional Scaling (MDS) was performed with R package “vegan” (version 2.5.4) at the species level and for AMR gene counts with the “Bray-Curtis” distance in the “capscale” function, and the p-values were extracted from the mixed linear models with effect of interests. Shannon diversity was calculated using the “diversity” function from the “vegan” package, and p-values were extracted from the two-sample t-test using the base R “t.test” function, or from Wilcoxon rank-sum test using the base R “wilcox.test” function. Mixed linear models were constructed using the “lme” function from the “nlme” package. Mixed linear models include subject ID as a random variable and patient group and time as fixed variables.

Statistical analyses and visualization of data was done through R version 4.2.2, and Python version 3.11.3.

## RESULTS

### Comparison of gut microbiomes in patients randomized to home versus hospital HCT demonstrates few differences between groups

Our study cohort was constructed from longitudinally sampled stools, from patients randomized to undergoing HCT either at home or in the hospital as part of Phase II trials, described previously^5^. We identified 28 patients who met inclusion criteria (see Methods); 16 of these were patients randomized to home HCT, 12 were randomized to hospital HCT. No significant differences were observed between groups for any patient- or transplant-related variables indicating appropriate randomization (**Table 1**). The duration of hospitalization was similar between the two groups (p=0.199; **Supplementary Figure S1**).

**Table 1.**
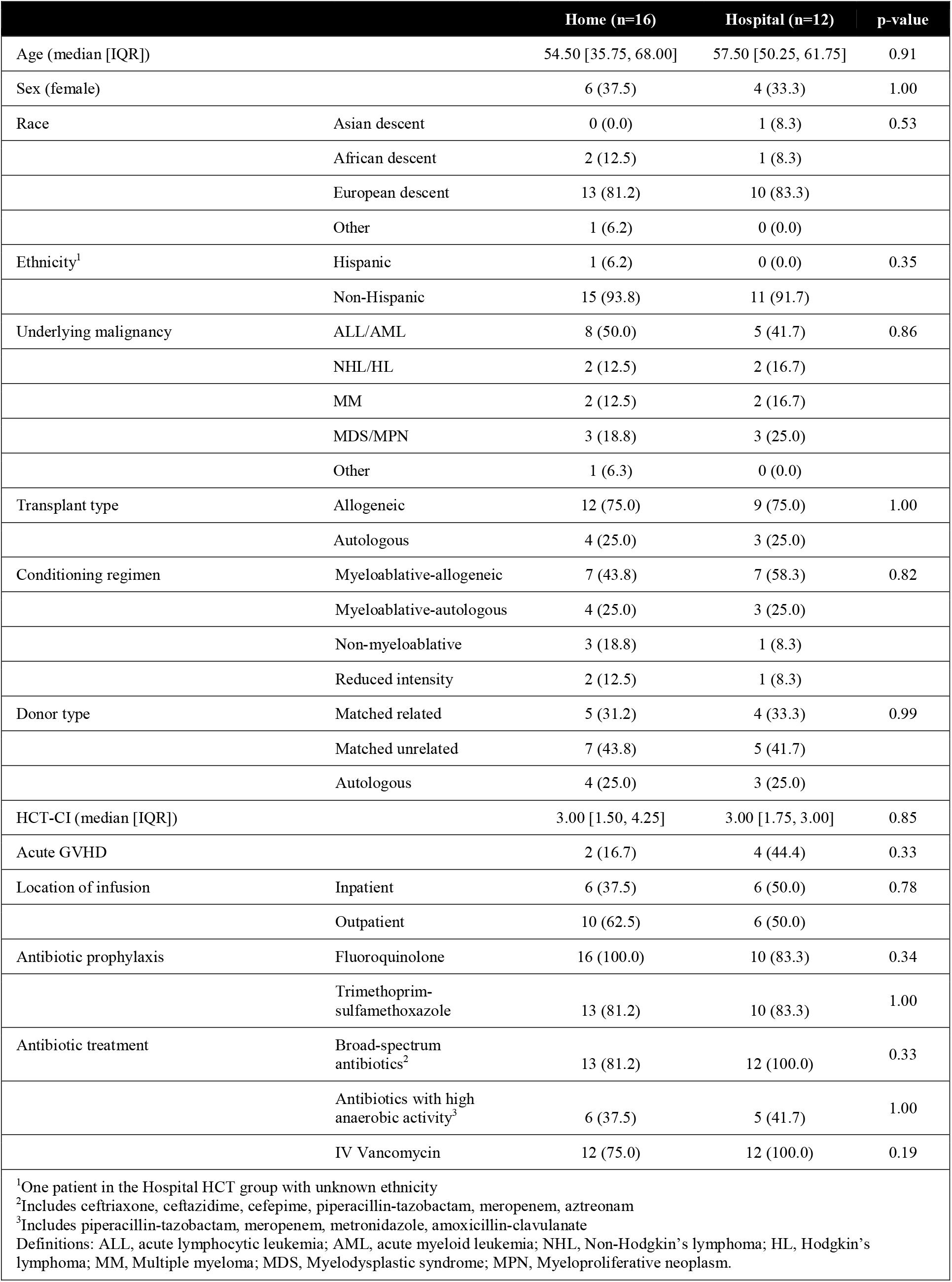
Demographics and transplant characteristics of patients included, comparing patients randomized to home and hospital HCT.

Stool samples from each patient were subjected to shotgun metagenomic sequencing to a target depth of 50-60 million 150 base pair paired-end reads. 213 stools were obtained, with only 203 stools sequenced as a result of low DNA concentrations following extraction for 10 samples. Stools ranged from -37 days prior to HCT to +767 after transplant (**Figure 1**). Sequenced reads were filtered to retain high-quality non-duplicated reads and human sequences were removed. Reads underwent both taxonomic characterization as well as characterization of antimicrobial resistance (AMR) genes via VSEARCH against the Comprehensive Antibiotic Resistance Database (CARD). Reads were assembled into contigs and orthogonal methods for AMR gene identification were applied: contigs were aligned against the CARD database with high stringency via RGI, and against NCBI using AMR Finder. On average, each sample was found to have a median of 3224 species (range 236-6249), while the median number of AMR genes identified varied widely between methods: 9 on average using AMR Finder (range 1-12), 54 using RGI (range 13-184), and 253 using VSEARCH (range 92-492).

**Figure 1.**
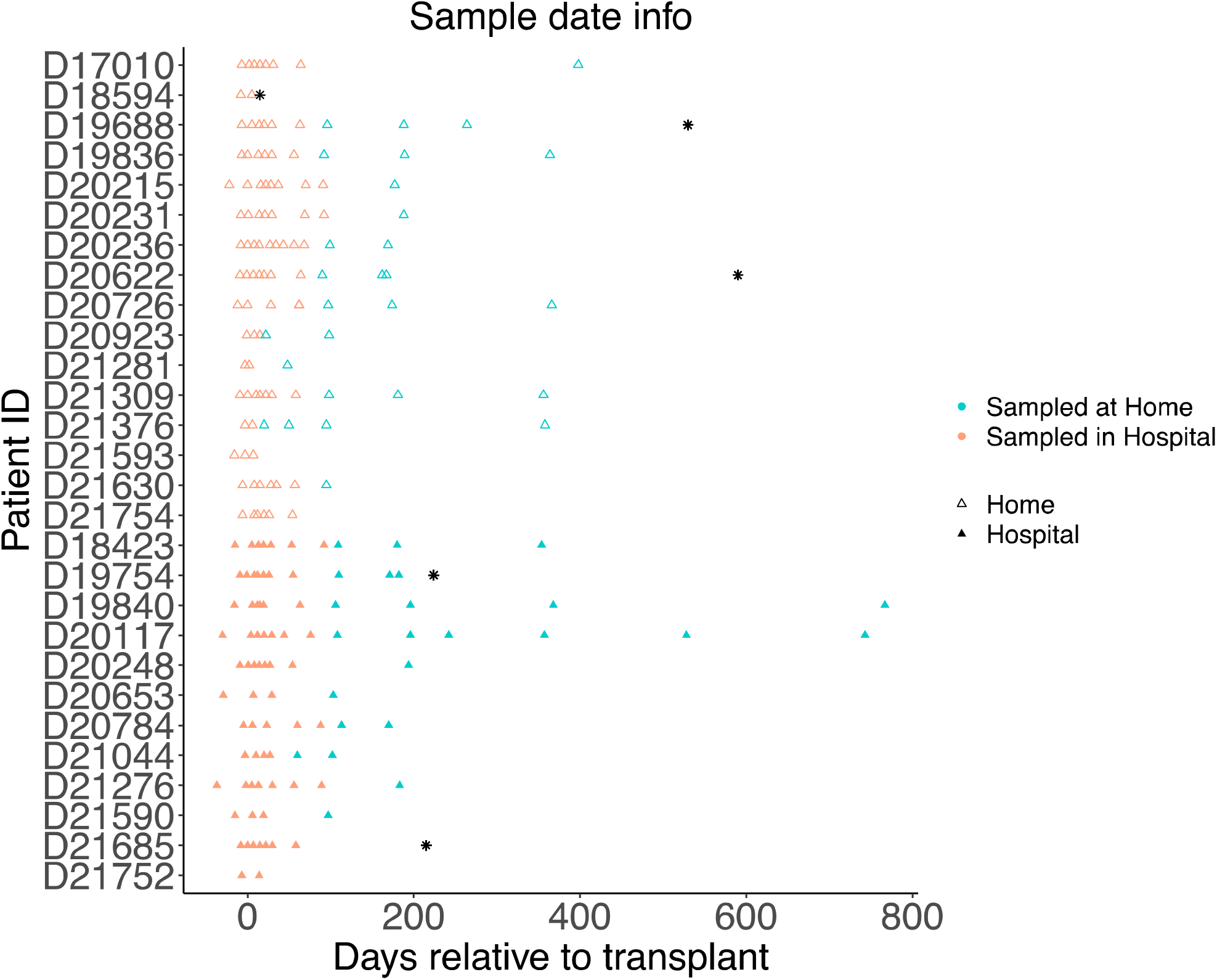
Stool samples from patients randomized to HCT at home or in the hospital are plotted for each patient relative to the day of transplantation. Samples are colored based on the location of sampling, shaped based on treatment group type; black asterisks represent the days after transplant on which patients died.

To compare taxonomic and AMR gene composition between patients who had undergone HCT at home versus in the hospital, we calculated Shannon diversity (**Figure 2)** before HCT (pre-HCT) and at days +30 and +60 after HCT at the species level (**Figure 2A**), for all timepoints (**Figure 2B**) and for AMRFinder, RGI, and VSEARCH results (**Figure 2C**). Overall, we find no significant difference in taxonomic Shannon diversity between home and hospital prior to or after transplantation, and no significant difference in AMR gene diversity at any time. No significant differences in home v. hospital HCT patients were observed when these same comparisons of Shannon diversity were performed separately for allo- and auto-HCT (data not shown).

**Figure 2.**
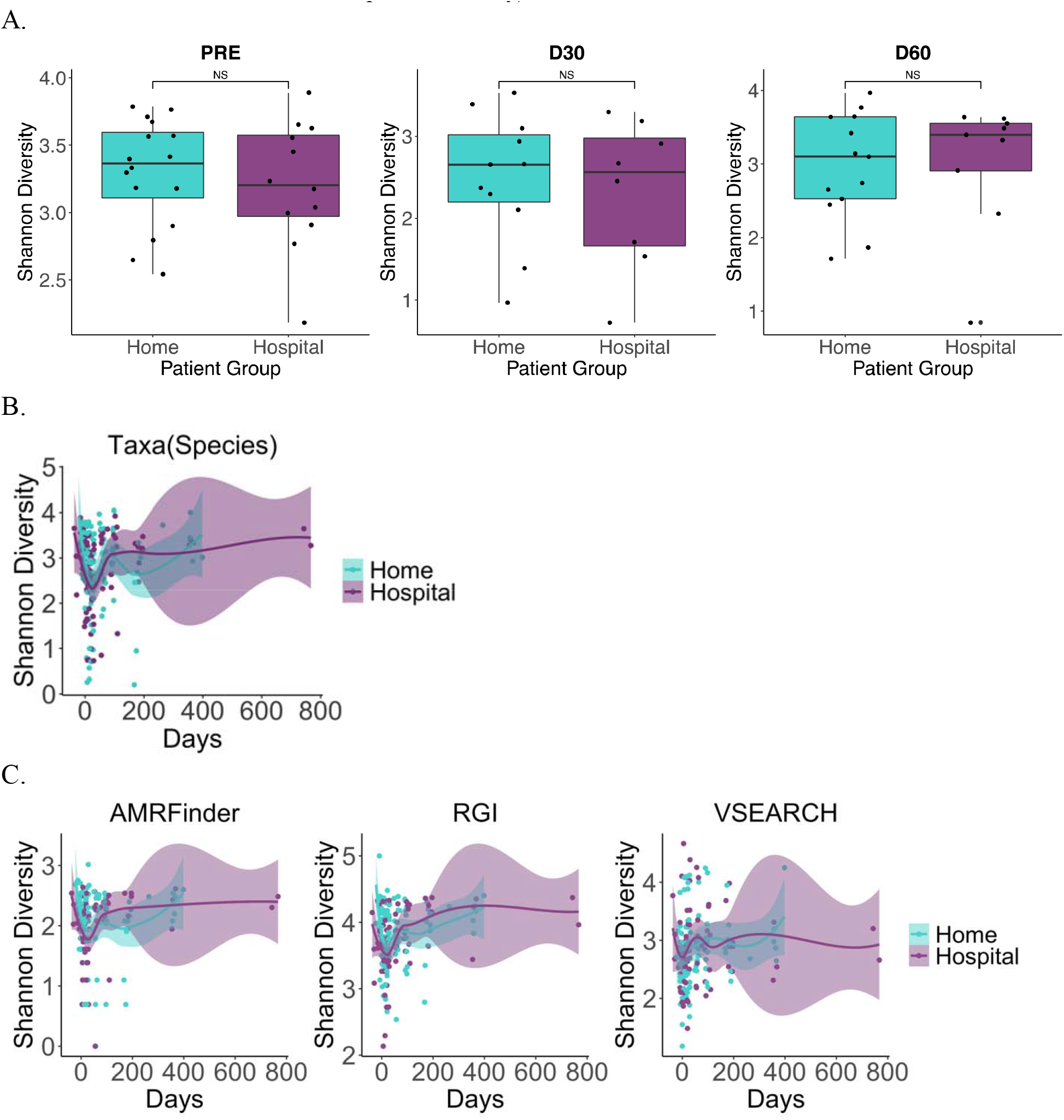
Taxonomic and AMR gene Shannon diversity did not differ between patients randomized to HCT at home or in the hospital. A) Taxonomic Shannon diversity at the species level was compared between patients undergoing home v. hospital HCT. No differences in Shannon diversity were found using the Wilcoxon rank-sum test before (p = 0.599) or after HCT at days +30 and +60 after transplant (p = 0.840 and p = 0.948, respectively). B) Shannon diversity over time at the species level is comparable between HCT groups (p = 0.519, mixed linear model). C) Shannon diversity of AMR genes characterized using three orthologous methods were not significantly different (AMRFinder, p = 0.351; RGI, p=0.177; VSEARCH, p=0.634, mixed linear model with home vs. hospital effect only).

In order to investigate taxonomic compositional, or beta-diversity, changes over time between home and hospital HCT groups, we compared Bray-Curtis dissimilarity between timepoints using multidimensional scaling (MDS); we plotted MDS1 for both taxonomic and AMR gene composition over time (**Figure 3**). For taxonomic composition and AMR genes, we observe no difference between groups at any time before or after transplant. No significant differences were observed when MDS was plotted separately for allo- and auto-HCT (data not shown).

**Figure 3.**
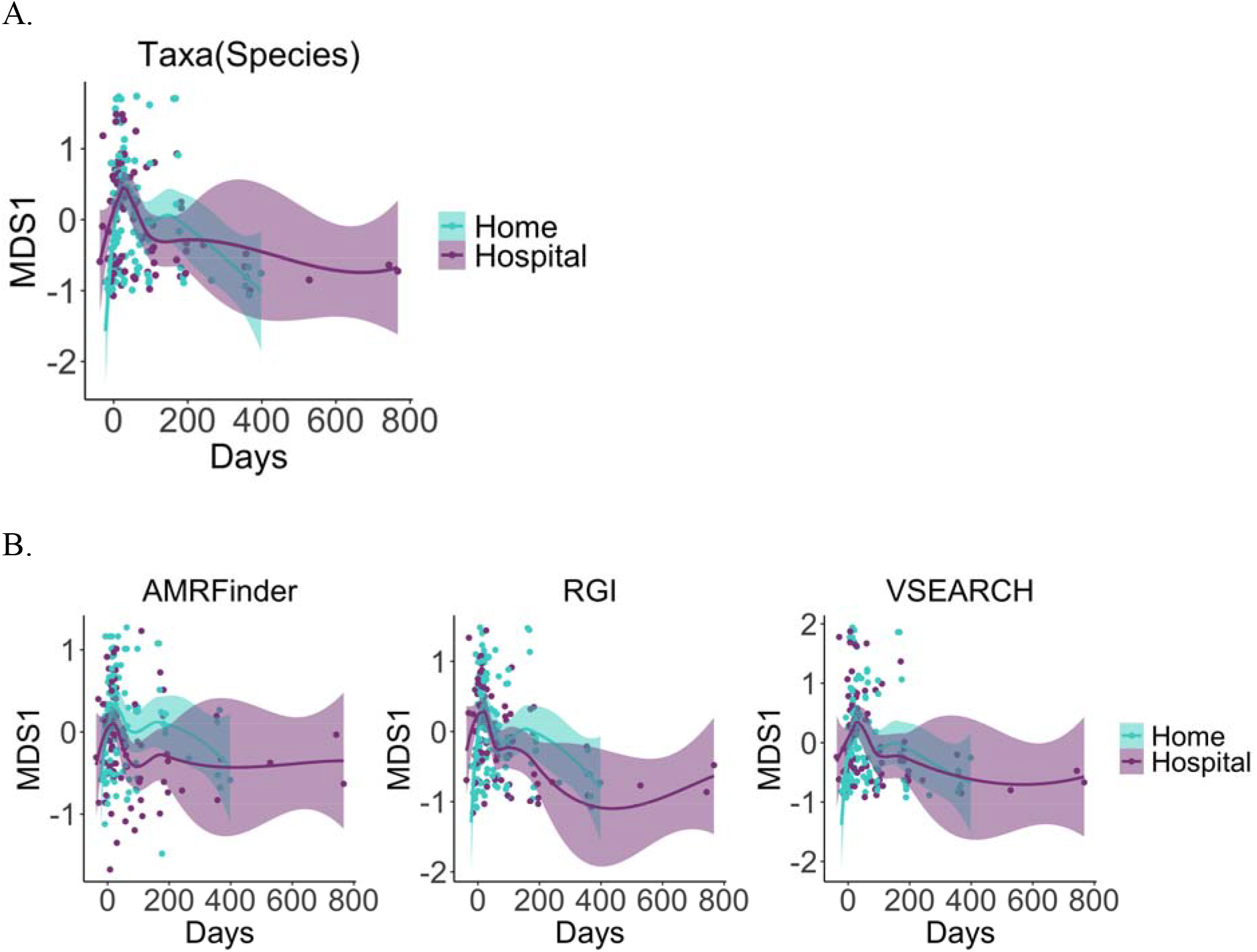
Beta-diversity of gut microbial taxonomic and AMR gene composition does not differ between home a d hospital HCT. Multidimensional scaling (MDS) between patients in each group at all timepoints was performed using Bray-Curtis dissimilarity measurements and tested via mixed linear models of A) gut microbiome taxonomic composition at the species level (p=0.482), B) and AMR genes characterized using three orthologous methods (AMRFinder, p = 0.155; RGI, p = 0.693; VSEARCH, p = 0.668).

We constructed mixed linear models in which randomization allocation (home v. hospital) and timepoint are fixed terms and subject ID is a random term. We ran these models on individual taxa, AMR genes, and metabolic pathways and found no statistically significant differences between groups after correction for multiple hypothesis testing (**Supplementary Figure 2**). However, we did observe changes in the differential abundance of *Streptococcus* spp. and *Actinomyces neuslundii* over time that trended towards significance (adjusted p-values=0.0532) with the relative abundance of these species declining or remaining stable in the home HCT group and rising in the hospital HCT group. We performed similar mixed linear models with AMR genes and metabolic pathways and did not find any significant differences between patients randomized to receive HCT at home or in the hospital (data not shown).

In summary, in a subgroup of patients from Phase II trials investigating the impact of transplant location on the gut microbiome, we find that the duration of hospitalization is similar between groups despite randomization to home or hospital HCT. Although we hypothesized that the α-diversity of patients randomized to home HCT would be lower than those in the hospital because of less regular intensive contact with the healthcare system and healthcare providers, we find no difference in either alpha- or beta-diversity, or any significant differences in taxa or AMR genes between groups.

### Secondary analyses demonstrate differences in gut microbiomes based on the location where patients spent most of their time after HCT, irrespective of randomized group allocation

Having observed no difference in the duration of hospitalization between patients randomized to home or hospital HCT, we sought to compare patients based on whether they spent more time in the inpatient or outpatient setting (see methods for further details on inpatient v. outpatient cohort definitions; **Supplementary Figure S3**). We found that the duration of hospitalization was significantly greater by 2 weeks for those patients allocated to receive their stem cell infusion in the inpatient setting (“inpatient”) versus those who received their infusion in the outpatient infusion clinic setting (“outpatient”) (29 days v. 5 days, p=5.41e-05; **Supplementary Figure S4**). We also found that in our cohort, the duration of hospitalization was inversely correlated with Shannon diversity for all patients (**Figure 4A**) and when considering only allo-HCT (**Figure 4B**). As the actual transplant location almost perfectly differentiated high versus low duration of hospitalization, in our secondary analysis we compared the gut microbiomes of these two groups (outpatient v. inpatient HCT) in order to determine the impact of hospitalization duration on gut microbial diversity.

**Figure 4.**
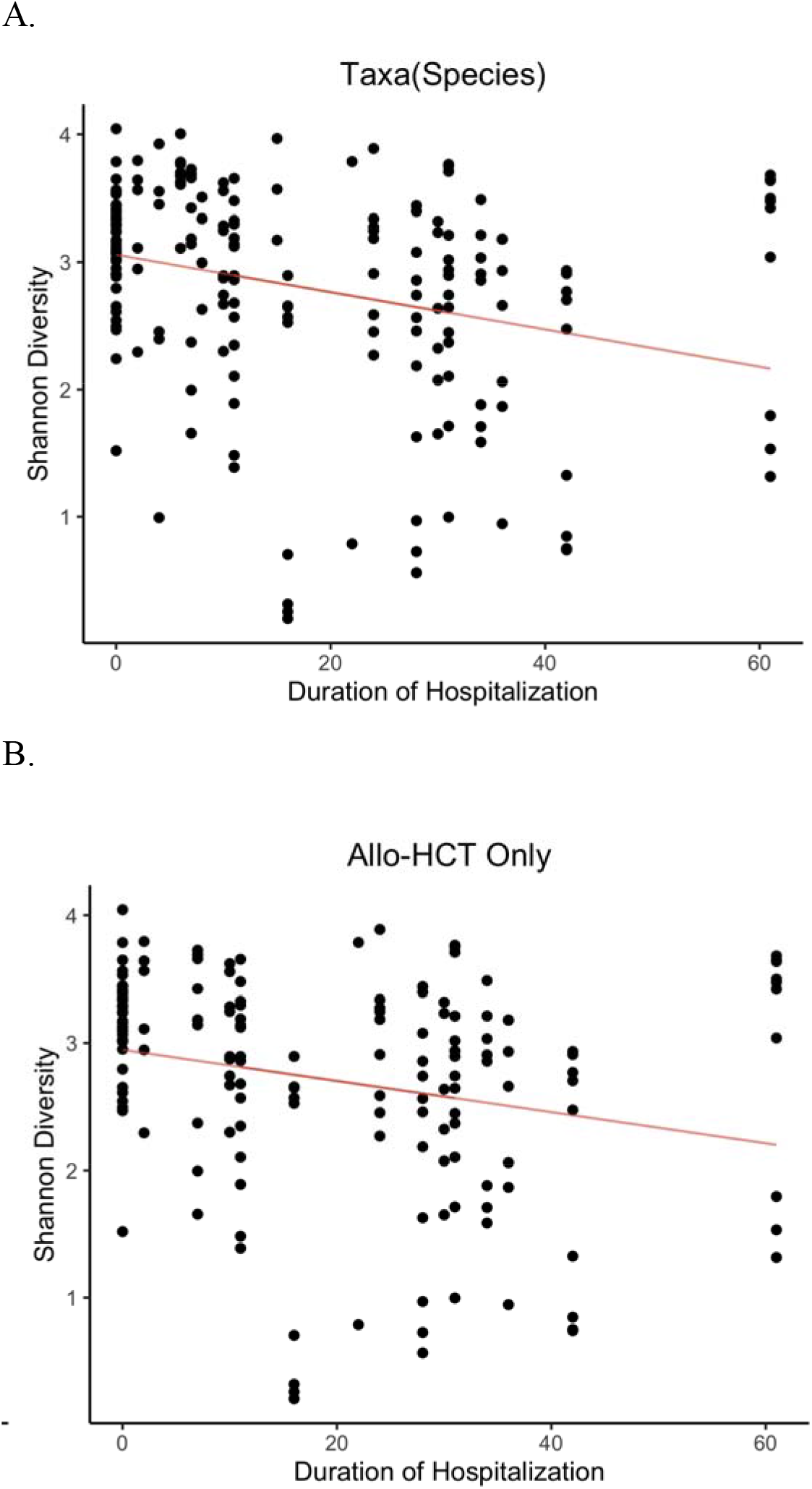
Shannon diversity was significantly lower in patients hospitalized for a longer duration. A) Taxonomic Shannon diversity at the species level was compared with hospitalization duration. Patients hospitalized for a longer duration were found to have significantly lower gut species Shannon diversity: A) when comparing all patients’ samples (p=0.00976, mixed linear model), and B) only including those patients undergoing allo-HCT (p=0.0439, mixed linear model).

In contrast to those undergoing home v. hospital HCT, patient and transplant variables between the outpatient and inpatient HCT groups were imbalanced (**Supplementary Table S1**). No differences were found between groups in patients’ age, race, or randomization to home v. hospital HCT groups. There were, however, significant differences in conditioning regimen and transplant type between groups with only samples from allogeneic recipients being obtained from the inpatient hospital group (p=0.03). Although the total number of antibiotic courses were not different between the two groups, patients in the inpatient HCT group were significantly more likely to receive antibiotics with higher anaerobic activity (*e*.*g*., piperacillin-tazobactam, meropenem) compared to those in the outpatient HCT group (p=0.003).

Comparing the gut microbiota between outpatient and inpatient HCT groups, we find no significant difference in Shannon diversity prior to transplantation, although outpatient HCT was significantly associated with higher diversity at both days +30 and +60 (**Figure 5A**; p < 0.05 by Wilcoxon test), and for all timepoints (**Figure 5B;** p < 0.05 by mixed linear model). In contrast, despite differential taxonomic diversity following transplant, no significant difference in AMR gene diversity was observed between groups before or after HCT (**Figure 5C**). As the two study groups were imbalanced in transplant type (*i*.*e*. only allogeneic HCT in the inpatient HCT group), we investigated differences in Shannon diversity throughout allo-HCT. When we compared only allo-HCT recipients, Shannon diversity remained significantly higher in the outpatient HCT group (**Figure 5B**).

**Figure 5.**
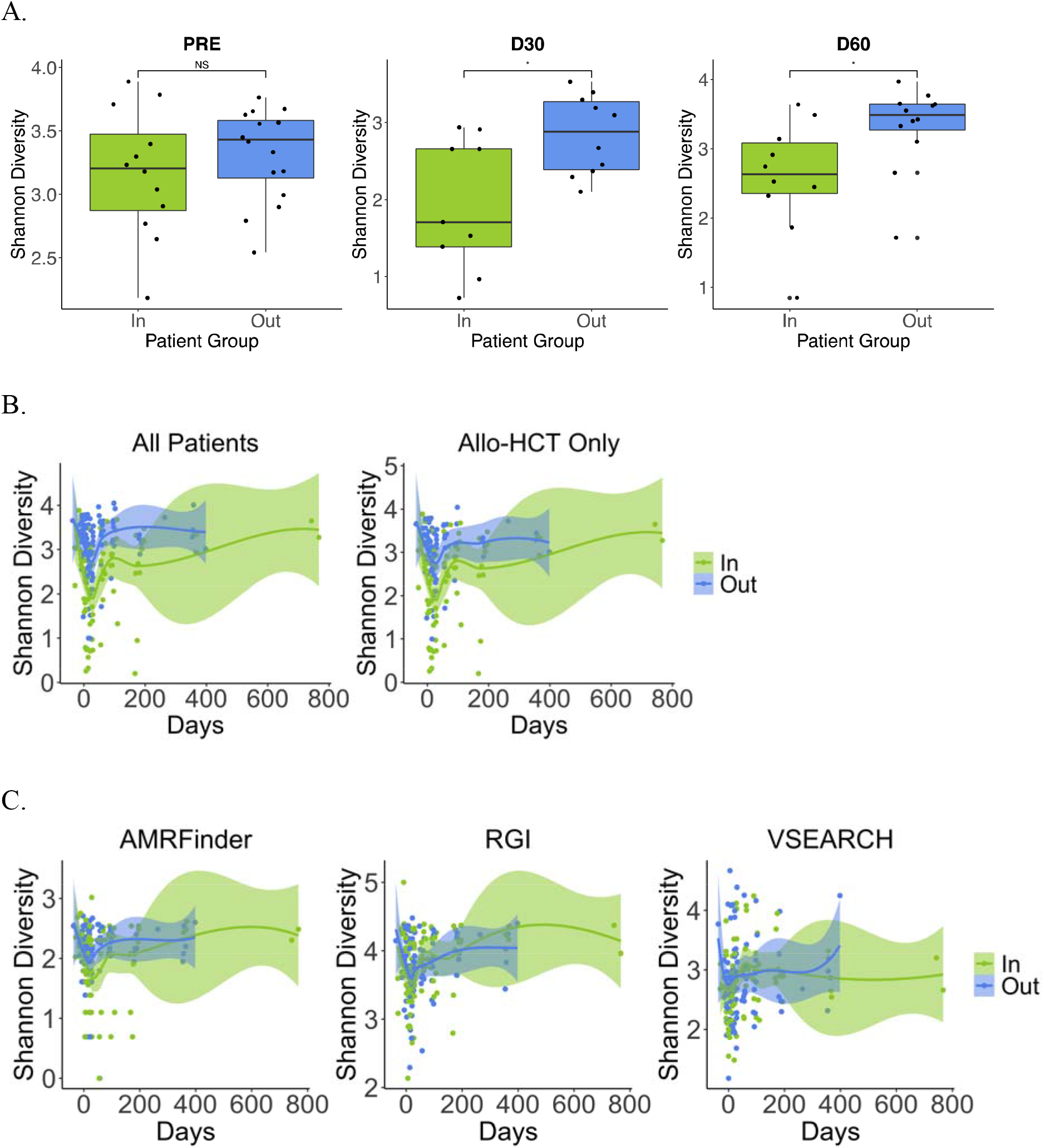
Shannon diversity was significantly higher in outpatient HCT recipients after, but not prior, to transplantation. A) Taxonomic Shannon diversity at the species level was compared between patients undergoing outpatient v. inpatient HCT. Patients in the outpatient HCT group were found to have higher gut species Shannon diversity at days +30 and +60 after transplant (p = 0.0279 and p = 0.0206 respectively) but not before HCT (p = 0.478, Wilcoxon rank-sum test). B) Shannon diversity over time at the species level is higher in the outpatient compared with the inpatient HCT group (p = 0.0000899, mixed linear model). The relative difference in taxonomic Shannon diversity between the groups remained significant even when only considering patients undergoing outpatient allo-HCT (n = 9) or inpatient allo-HCT (n = 12) (p = 0.00132, mixed linear model). C) Shannon diversity of AMR genes characterized using three orthologous methods were not significantly different (AMRFinder, p = 0.155; RGI, p = 0.645; VSEARCH, p = 0.984).

In terms of β-diversity, for both taxonomic composition and AMR genes, we observe little difference between groups pre-HCT but more separation between inpatient and outpatient HCT at after transplant (p = 0.00661, mixed linear model with inpatient vs. outpatient effect only) (**Figure 6**). We found that when comparing differential taxonomic composition between groups over time using MDS1, β-diversity was significantly higher in outpatient compared to inpatient HCT recipients (**Figure 6A**), although these same differences were not observed consistently for AMR genes and observed only in VSEARCH results (**Figure 6B**).

**Figure 6.**
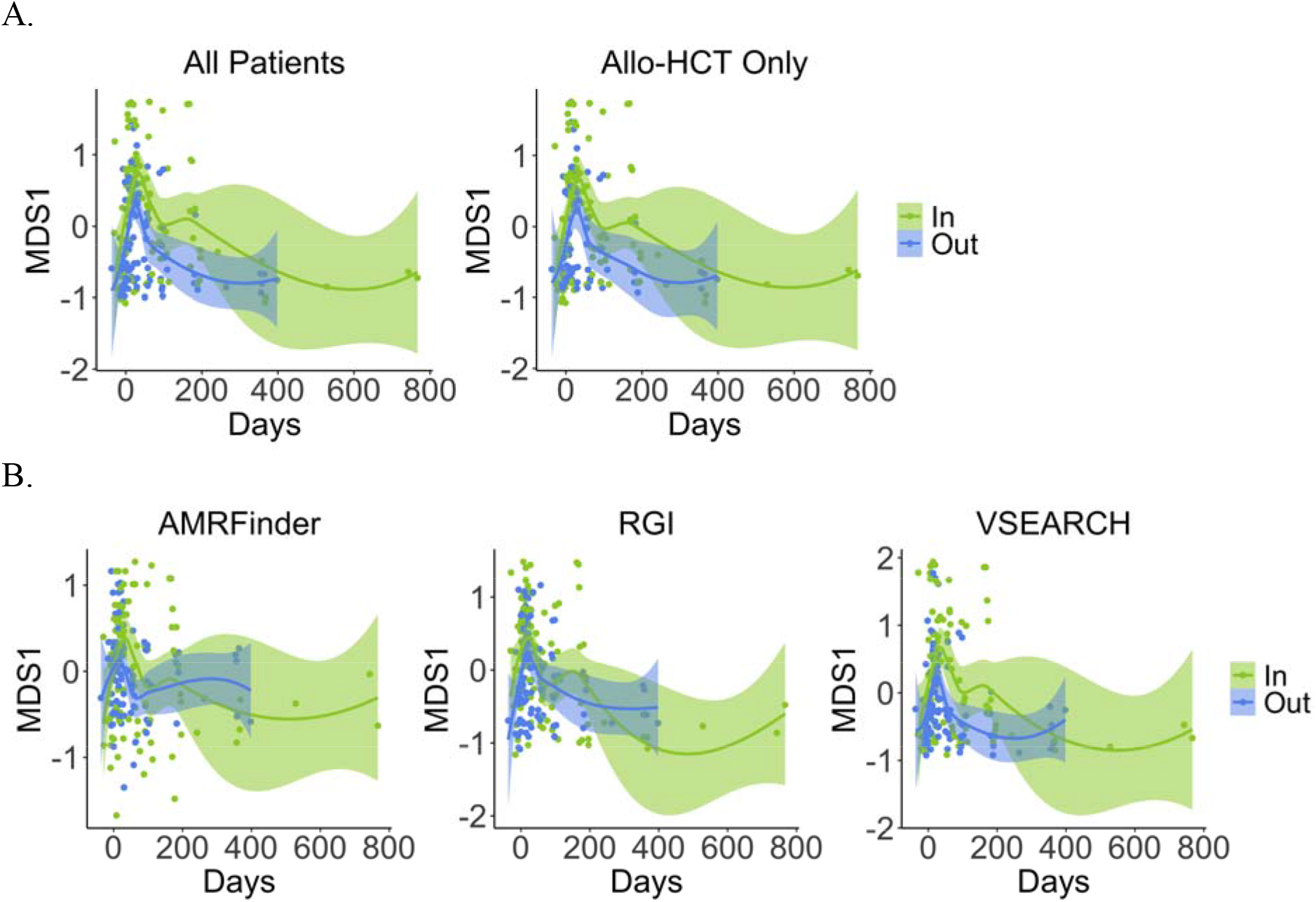
Beta-diversity of gut microbial taxonomic and AMR gene composition differs after, but not before, transplant between inpatient and outpatient HCT groups. Multidimensional scaling (MDS) between patients in each group at all timepoints was performed using Bray-Curtis dissimilarity measurements and testing via mixed linear models of A) gut microbiome taxonomic composition at the species level (p = 0.00661), when considering allo-HCT patient only (p = 0.0318), and B) AMR genes characterized using three orthologous methods (AMRFinder, p = 0.405; RGI, p = 0.110; VSEARCH, p = 0.0173).

We next constructed mixed linear models in which transplant location (outpatient vs. inpatient HCT) and timepoint are fixed terms and subject ID is a random term. After correction for multiple hypothesis testing, we did not find any significant differential abundance in individual taxa or AMR genes between groups over time (data not shown). These results suggest a modest effect size in beta-diversity for the relationship between transplant location and taxonomic composition, but not enough of an effect size that could be observed with individual taxa alone.

## DISCUSSION

Recent years have seen an increasing demand for home-based healthcare, recognizing the impact that prolonged hospitalization can have on patients’ quality of life. Trials of home versus hospital HCT have demonstrated that home-based HCT proves to be a safe alternative to inpatient treatment with benefits for patients’ overall wellbeing, however the impact that the shift from hospital to home environment and the effect of the built environment on the gut microbiome has not yet been established. The Karolinska Institute reported a decrease in acute GVHD incidence in their home HCT patients although these patients were not randomized^3^. Based on this, we hypothesized that these differences in GVHD outcomes between home and hospital HCT recipients may be due to gut microbial taxonomic differences between groups. In this first of its kind investigation, we demonstrate in patients randomized to home v. hospital HCT, that there were no significant differences in Shannon diversity or gut microbial taxonomic composition before or after transplant. Despite expected differences in the level of interaction with the health care system between the home vs. hospital groups, patients were hospitalized for a similar duration in those randomized to home or hospital HCT which may be one of the major reasons why their microbiomes were also similar. In a secondary analysis, we compared groups based on whether they underwent the majority of their transplant care in the inpatient or outpatient setting. Patients in the inpatient vs. outpatient groups had significant differences in overall duration of hospitalization. Inpatient HCT recipients were found to have a significantly lower Shannon diversity and differential taxonomic composition after, but not before, transplant compared to outpatient HCT recipients. These differences in groups were highly confounded however, including by the type of transplant with only allogeneic recipients in the inpatient HCT group. To address this, we compared only allogeneic HCT patients in gut diversity, composition, and Bray-Curtis dissimilarity from baseline, and found similar results, many of which were statistically significant despite the smaller group size; this rules out transplant type as a primary driver of our results. Apart from important confounders such as transplant type that require consideration, inpatient HCT recipients were far more likely to have received antibiotics with significant anaerobic activity compared to their outpatient HCT counterparts, although there were no differences in the frequency of broad-spectrum antibiotics overall (**Table S1**). Prior studies have demonstrated a significant impact of anaerobically active antibiotics on the gut microbiome and worse clinical outcomes in HCT recipients^24,25^.

Interestingly, differences in diversity and taxonomic composition based on actual transplant location, did not translate into differences in AMR gene diversity. While our analyses benefitted from an orthogonal approach to characterizing AMR genes using three methods, our analysis did not compare differences in acquired AMR genes. Future work investigating the impact of home transplant on patients’ microbiota will benefit from investigation of the “mobilome” of highly mobile antimicrobial resistance genes that may yield more evident differences in AMR gene acquisition based on duration of healthcare exposure. While we observed modest differences based on actual HCT location, these were confounded heavily by patient and transplant variables. Any impact of HCT location due to variability in diet and exposure may therefore be too difficult to untangle from the factors that significantly impact patients’ overall disease and transplant trajectory.

## Acknowledgements

We especially thank the patients and nurses on the Blood and Marrow Transplantation service at Duke University for their participation in this project. This work was supported in part by an Amy Strelzer Manasevit Award from the National Marrow Donor Program, an award from the National Institute of Allergy and Infectious Diseases (K23 AI163365), and the UNC Physician-Scientist Training Award (T.M.A.). A.D.S. and N.J.C. received awards for this work from NHLBI (R01AG066719 to A.D.S. and R01CA203950 to N.J.C.).

## Conflicts of Interest

The authors report no conflicts of interest.

## Notes

### Competing Interest Statement

The authors have declared no competing interest.

