## Supplemental Tables and Figures for "The impact of transplant location on the gut microbiome and resistome in patients undergoing hematopoietic stem cell transplantation at home versus in the hospital"

**SUPPLEMENTARY FIGURES AND TABLES**

**Figure S1. Similar duration of hospitalization observed for patients randomized to HCT in the hospital compared to the home.** We compared the duration of intensive daily transplant care using the Wilcoxon rank-sum test between patients randomized to HCT at home or in the hospital and found no significant difference between groups (p=0.199).

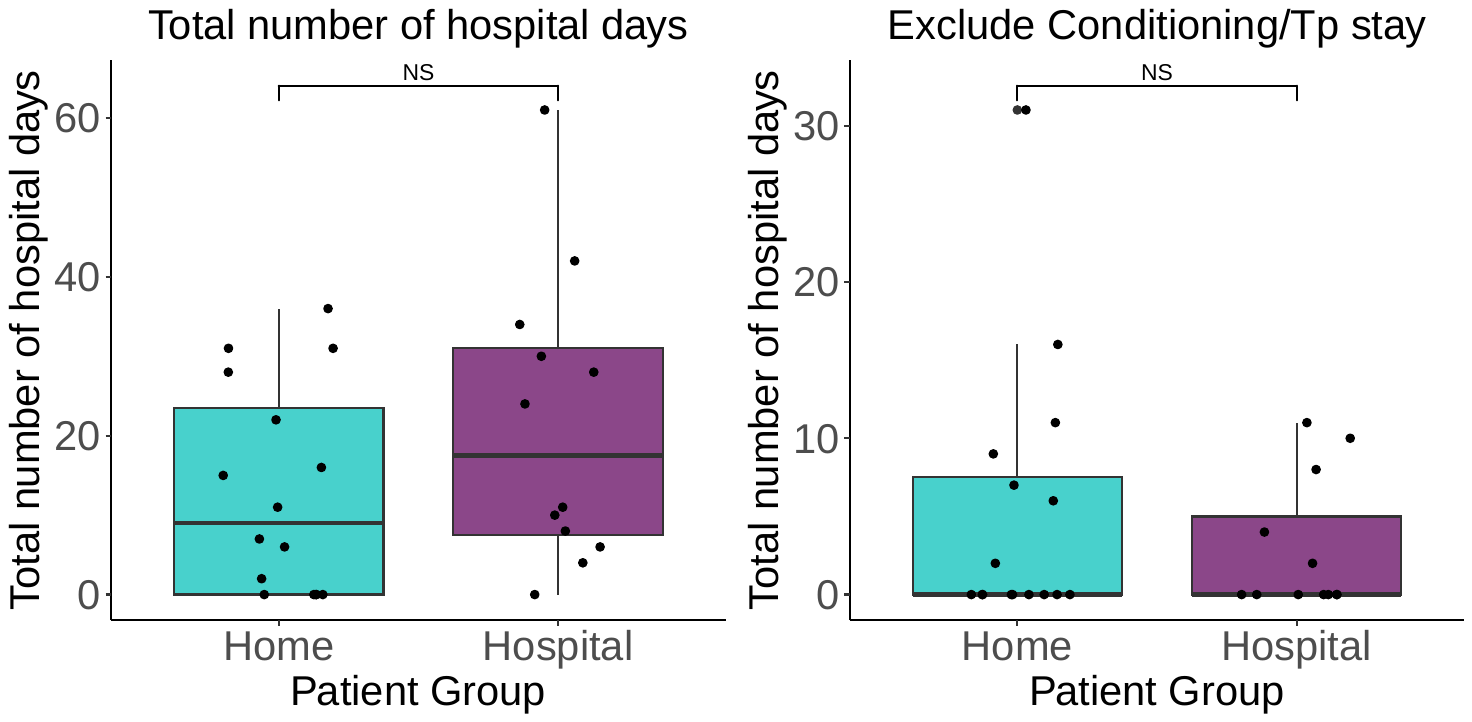

**Figure S2.** **Mixed linear models demonstrate no individual taxa or metabolic pathways that differ significantly between patients randomized to home or hospital HCT.** Top eight most significant taxa at the species level (p-values from mixed linear models with home vs. hospital effect only).

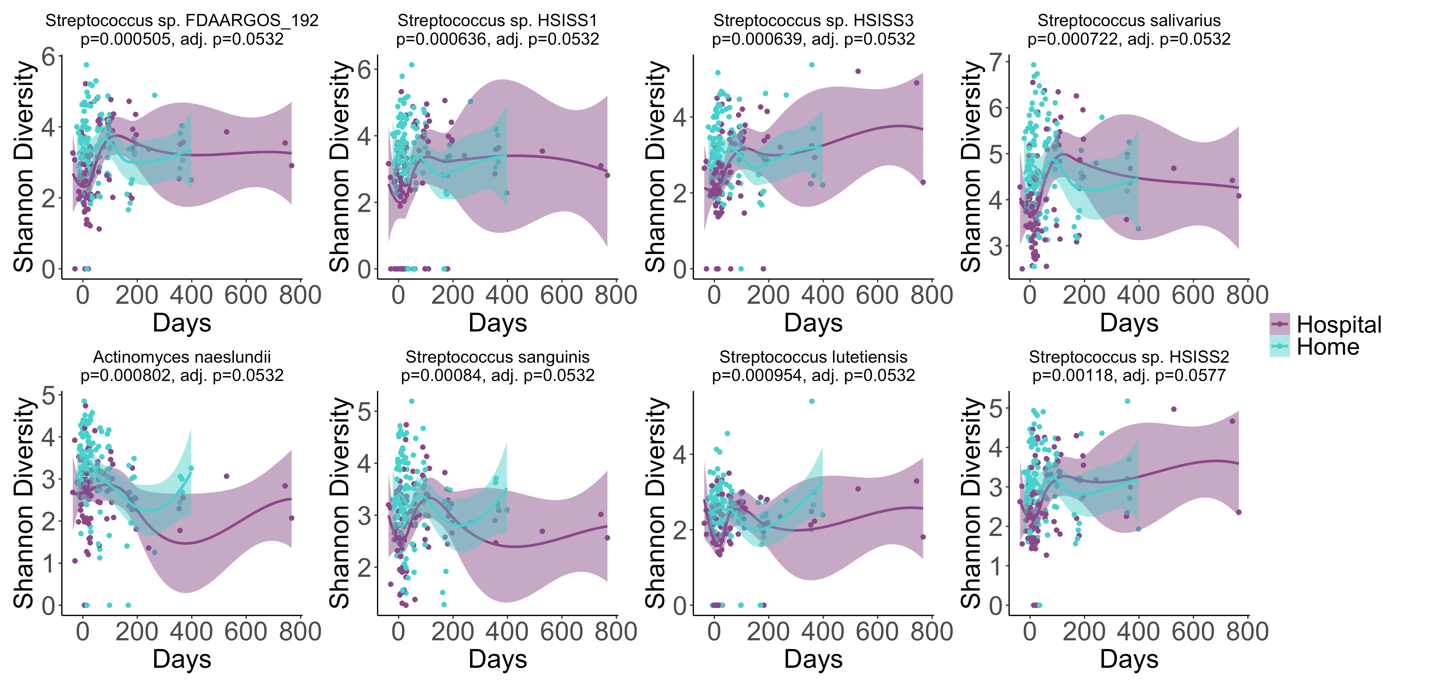

**Figure S3. Stool samples from patients undergoing inpatient versus oputpatient HCT are plotted for each patient relative to the day of transplantation.** Samples are colored based on the location of sampling, shaped based on treatment group type; black asterisks represent the days after transplant on which patients died.

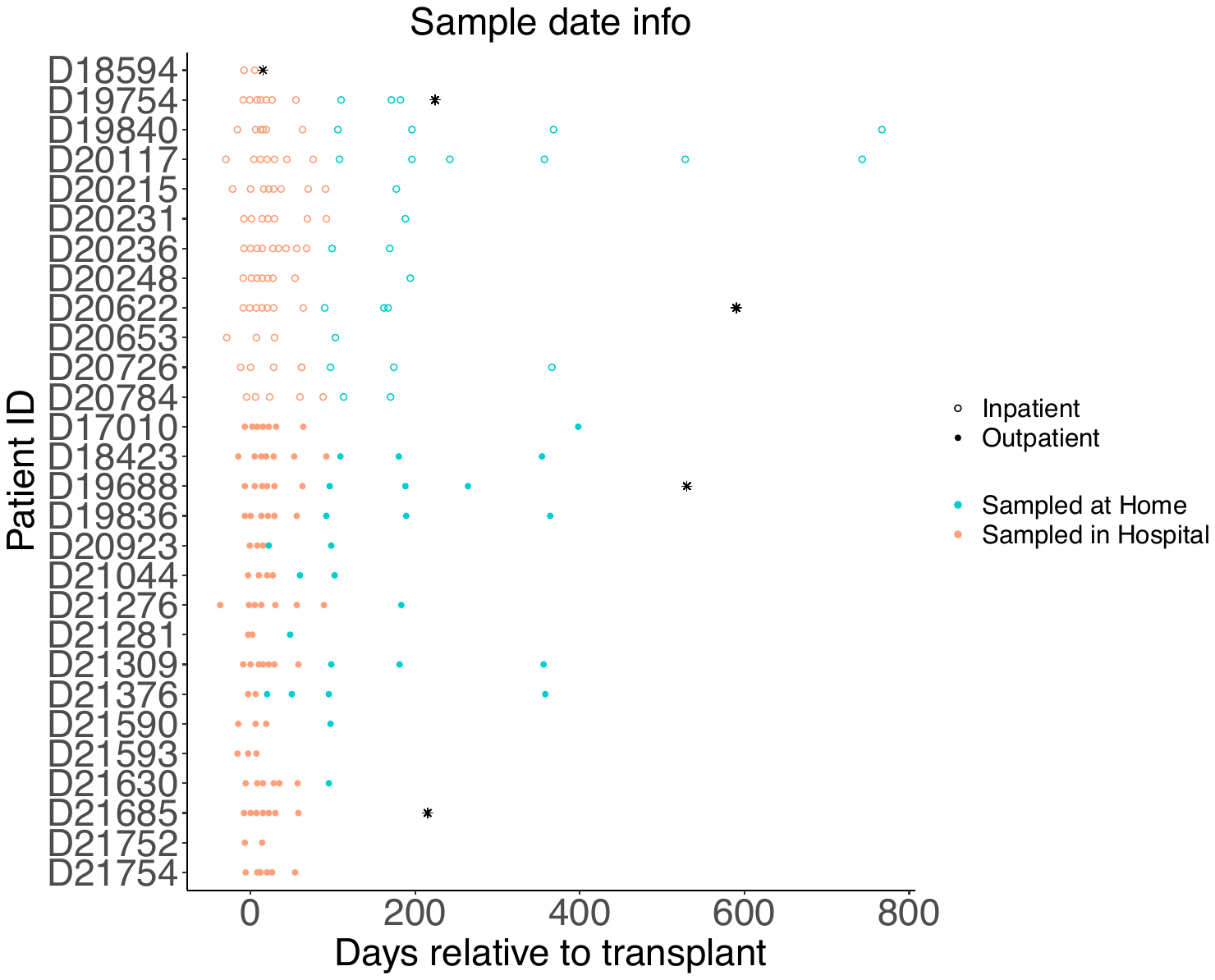

**Figure S4.**  **Total number of hospital days observed for patients in the inpatient HCT group is significantly higher compared to the outpatient HCT group.** We compared the duration of intensive daily transplant care using the Wilcoxon rank-sum test between patients assigned to HCT in the inpatient or outpatient settings by patient (p =5.41e-05).

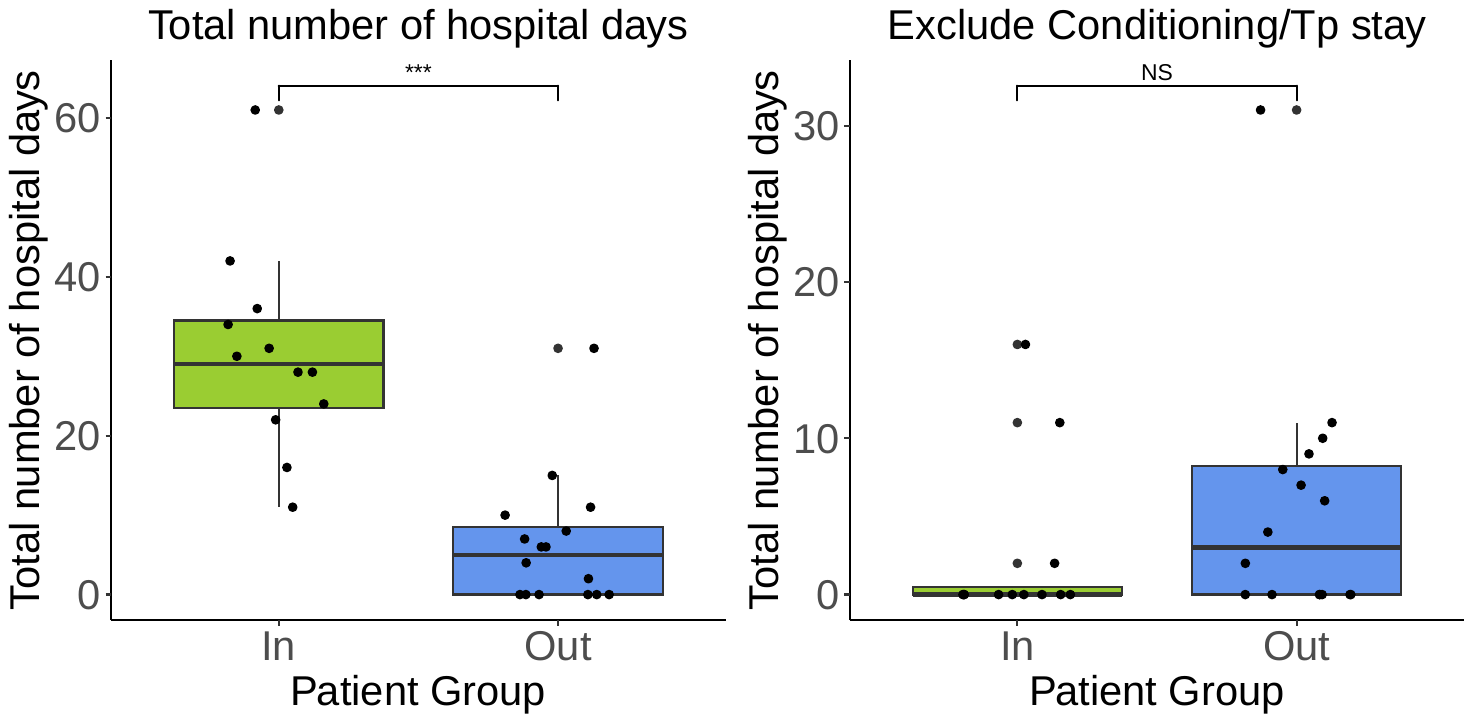

**Table S1.** Demographics and transplant characteristics of patients included, comparing patients undergoing outpatient and inpatient HCT.

|  |  | **Inpatient (n=12)** | **Outpatient (n=16)** | **p-value** |
| --- | --- | --- | --- | --- |
| Age (median [IQR]) |  | 52.00 [41.00, 60.25] | 60.50 [43.00, 67.25] | 0.46 |
| Sex (female) |  | 6 (50.0) | 4 (25.0) | 0.33 |
| Race | Asian descent | 1 (8.3) | 0 (0.0) | 0.41 |
|  | African descent | 2 (16.7) | 1 (6.2) |  |
|  | European descent | 9 (75.0) | 14 (87.5) |  |
|  | Other | 0 (0.0) | 1 (6.2) |  |
| Ethnicity^1^ | Hispanic | 1 (8.3) | 0 (0.0) | 0.35 |
|  | Non-Hispanic | 11 (91.7) | 15 (93.8) |  |
| Underlying malignancy | ALL/AML | 8 (66.7) | 5 (31.3) | **0.02** |
|  | NHL/HL | 0 (0) | 4 (25.0) |  |
|  | MM | 0 (0) | 4 (25.0) |  |
|  | MDS/MPN | 4 (33.3) | 2 (12.5) |  |
|  | Other | 0 (0) | 1 (6.3) |  |
| Transplant type | Allogeneic | 12 (100.0) | 9 (56.2) | **0.03** |
|  | Autologous | 0 (0.0) | 7 (43.8) |  |
| Conditioning regimen | Myeloablative-allogeniec | 11 (91.7) | 3 (18.8) |  |
|  | Myeloablative-autologous | 0 (0.0) | 7 (43.8) | **0.002** |
|  | Non-myeloablative | 1 (8.3) | 3 (18.8) |  |
|  | Reduced intensity | 0 (0.0) | 3 (18.8) |  |
| Donor type | Matched related | 4 (33.3) | 5 (31.2) | **0.02** |
|  | Matched unrelated | 8 (66.7) | 4 (25.0) |  |
|  | Autologous | 0 (0) | 8 (47.1) |  |
| HCT-CI (median [IQR]) |  | 2.50 [0.75, 3.00] | 3.00 [2.00, 4.25] | 0.24 |
| Acute GVHD |  | 5 (41.7) | 1 (11.1) | 0.18 |
| Randomized to home HCT |  | 6 (50.0) | 10 (62.5) | 0.78 |
| Antibiotic prophylaxis | Fluoroquinolone | 12 (100.0) | 14 (87.5) | 0.596 |
|  | Trimethoprim-sulfamethoxazole | 11 (91.7) | 12 (75.0) | 0.522 |
| Antibiotic treatment | Broad-spectrum antibiotics^2^ | 12 (100.0) | 13 (81.2) | 0.332 |
|  | Antibiotics with high anaerobic activity^3^ | 9 (75.0) | 2 (12.5) | **0.003** |
|  | IV Vancomycin | 12 (100.0) | 12 (75.0) | 0.185 |
| ^1^One patient in the Home HCT group with unknown ethnicity  ^2^Includes ceftriaxone, ceftazidime, cefepime, piperacillin-tazobactam, meropenem, aztreonam  ^3^Includes piperacillin-tazobactam, meropenem, metronidazole, amoxicillin-clavulanate  Definitions: ALL, acute lymphocytic leukemia; AML, acute myeloid leukemia; NHL; MM; MDS; MPN. | | | | |
